# Hippocampus-evoked polysynaptic responses in the medial prefrontal cortex are attenuated in aged rats

**DOI:** 10.64898/2026.06.05.730244

**Authors:** Sahana Srivathsa, Abhilasha Vishwanath, Victoria Garza, Stephen L. Cowen, Carol A. Barnes

## Abstract

The hippocampus-medial prefrontal cortex (mPFC) circuit is critical for spatial working-memory and decision-making. Age-associated changes to these functions are attributed to alterations in this circuit, although the exact mechanisms remain unclear. These regions are connected via monosynaptic projections from the intermediate (iHC) and ventral (vHC) hippocampus to the mPFC. We examined the functional connection between these regions in young (10-12 months) and aged (23-26 months) male F344 rats. High-density Neuropixels probes were used to record mPFC field excitatory postsynaptic potentials (fEPSPs) and single-unit responses to electrical stimulation of iHC and vHC. We observed that hippocampal stimulation evoked both short-latency (5-35 ms) monosynaptic and long-latency (35-100 ms) polysynaptic responses across the dorsoventral mPFC axis. Monosynaptic fEPSP amplitude increased with mPFC depth for iHC and vHC stimulation sites. Age did not affect hippocampal-prefrontal monosynaptic transmission as measured by fEPSP amplitude and evoked neural firing. In contrast, polysynaptic responses, reflecting local mPFC circuit recruitment, were larger than monosynaptic responses in young but not aged rats. Polysynaptic fEPSPs increased toward ventral mPFC in young but not aged rats, and polysynaptic evoked firing was reduced with age, most prominently at ventral mPFC depths. Our findings demonstrate that aging preserves the functional direct synaptic transmission but impairs polysynaptic recruitment of local mPFC circuits, with ventral mPFC showing particular vulnerability. These circuit-level deficits may contribute to age-related impairments in cognitive flexibility.

**Significance Statement:** Aging impairs spatial working-memory and decision making, which require hippocampus-prefrontal communication. We examined age-related changes in the functional connections of direct projections from the intermediate and ventral hippocampus onto dorsal and ventral regions of the mPFC. Direct monosynaptic HC-mPFC transmission was preserved with age. In contrast, evoked polysynaptic excitatory field potentials and single-unit responses, reflecting local mPFC circuit recruitment, were attenuated in aged rats, most prominently at ventral mPFC depths, and these changes differed between HC stimulation sites. These results identify subregion-specific reductions in the recruitment of local mPFC circuits by hippocampal input in aging, which may contribute to age-related impairments in cognitive flexibility.

## Introduction

Aging negatively impacts episodic and working-memory, as well as executive functions that rely on them (Barnes, 1979; Bizon et al., 2012; Wang et al., 2011; Kapellusch et al., 2018; Sotoudeh et al., 2020; Caetano et al., 2012). These forms of memory are supported by interactions between the hippocampus and medial prefrontal cortex (mPFC) (Floresco et al., 1997; Baddeley et al., 2000; Wang & Cai, 2006; Churchwell et al., 2010; Sapiurka et al., 2016; Ito, 2018). It is believed that the hippocampus delivers contextual information to the mPFC, supporting the transfer and integration of spatial and contextual representations to guide flexible behaviors (Baddeley et al., 2000; Moscovitch et al., 2005; Churchwell & Kesner, 2011; Preston & Eichenbaum, 2013; Anderson & Floresco, 2022; Moscovitch et al., 2016). Neural interactions between the dorsal hippocampus (dHC) and mPFC in rodents have been well characterized in spatial working-memory and decision-making tasks (Lee & Kesner, 2003; Jones & Wilson, 2005; Churchwell et al., 2010; Hyman et al., 2010; Jadhav et al., 2016; Griffin, 2015; Shin et al., 2019; Zielinski et al., 2019; Yadav et al., 2022; Wirt & Hyman, 2017; Nardin et al., 2023; Tang et al., 2017, 2023); however, these regions are only indirectly connected via bidirectional connections through the thalamus (Vertes, 2006; Hoover & Vertes, 2007; Varela et al., 2014; Griffin, 2021). A unidirectional projection from the intermediate and ventral CA1 region of hippocampus (iHC, vHC) to the mPFC directly connects these regions (Jay & Witter, 1991; Hoover & Vertes, 2007). Interactions of this iHC/vHC-mPFC circuit are primarily associated with context-dependent anxiogenic behaviors (Bannerman et al., 2003; Maren & Holt, 2004; Adhikari et al., 2010; Jin & Maren, 2015; Ciocchi et al., 2015; Padilla-Coreano et al., 2016; Q. Wang et al., 2016; Phillips et al., 2019; Marek et al., 2018; Sapiurka et al., 2016), but have also been known to influence spatial working-memory (O’Neill et al., 2013; Benchenane et al., 2010; Spellman et al., 2015; Tamura et al., 2017; Park et al., 2021; Dickson et al., 2022; Schoenfeld et al., 2023; Babl & Sigurdsson, 2025). The deterioration of spatial working-memory with age may therefore be explained by changes in the functional connectivity of the iHC/vHC input to the mPFC. Although no rodent work has yet explored how aging alters the hippocampal-prefrontal pathway, human neuroimaging studies suggest that both the functional connectivity and white matter tracts between the HC and PFC are reduced with age, correlating with changes in episodic memory (Ankudowich et al., 2019; Gunbey et al., 2014; Metzler-Baddeley et al., 2011; Stadlbauer et al., 2008).

Within the mPFC, the dorsal-prelimbic and ventral-infralimbic cortex regions receive direct monosynaptic glutamatergic projections from the iHC and vHC (Jay et al., 1992; Hoover & Vertes, 2007) although the density of these projections differs by depth and HC stimulation site (Sánchez-Bellot et al., 2022; Anastasiades & Carter, 2021; Alemán-Andrade et al., 2025). Recent anatomical tracing and slice physiology studies reveal vHC preferentially innervates neurons in the ventral mPFC regions, with fewer projections to dorsal mPFC regions (Liu & Carter, 2018; Little & Carter, 2012, 2013; Hoover & Vertes, 2007; Alemán-Andrade et al., 2025). Here, we assess the impact of age on the HC-mPFC circuit using large-scale *in vivo* recordings to compare evoked field potential and single-unit responses to iHC/vHC stimulation across the dorsoventral gradient of the mPFC. We examine how age affects the magnitude and latency of monosynaptic and polysynaptic responses in young and old rats to determine whether there are changes in HC-mPFC synaptic strength or circuit dynamics that could explain age-related deficits in spatial working-memory.

## Methods

### Subjects

Six young (10-12 months) and six old (23-26 months) male Fischer 344 rats (275-341 g) were used for this experiment. All animals were individually housed, kept on a reversed light/dark cycle (12 hours each) and fed *ad libitum* prior to the experiment. One young and one old rat were excluded from the analysis due to noise in the neural recordings which prevented reliable detection of spikes. This resulted in a total of n=5 rats used in the analysis per age group. Prior to the experiment, spatial memory on the Morris watermaze task was assessed for 3 out of 5 rats in each age group to verify an age deficit in spatial memory as previously shown (Gage et al., 1984).

### Surgery

Rats were anesthetized with isoflurane (VetOne, VGP Group LLC) in oxygen (1.5 L/min), induced at 4%, and maintained at 0.8-1.2% throughout surgery. Anesthesia dose was monitored by respiratory rate using an automated respiration-rate detector (38-42 breaths/minute; RespiRat, University of Arizona) and tail-pinch withdrawal was tested every 15 minutes. Core temperature was maintained with a water-circulating heating pad (T/Pump, Stryker). A bipolar concentric stimulating electrode (Plastics One MS303/1, 1 mm pole separation) was placed into the right hippocampus (centered at AP: 5.8 mm, ML: 5.4 mm from bregma), with depth adjusted for intermediate (3.5 mm below dura) or ventral (6.8 mm below dura) hippocampus stimulation (**Figure 1A**). A four-shank Neuropixels 2.0 probe (Steinmetz et al., 2021) was placed ipsilaterally in mPFC (AP: +3.0 mm, ML: 0.5-0.7 mm). The stimulation electrode and recording probes were initially lowered 0.1 mm beyond their target and then slowly retracted to minimize tissue drift. Data were recorded from 384 recording sites on the probe (2.175 - 5.025 mm below dura) along all 4 shanks (separated by 0.25mm, spanning 0.5-1.5 mm laterally; **Figure 1B-D**). An Ag/AgCl reference electrode (0.25 mm diameter silver wire) was prepared by soaking silver wire in 8.25% sodium hypochlorite and placing it in the ipsilateral cortex (AP: 3.0 mm, ML: 2.5 mm, DV: 1.0 - 1.2 mm). Craniotomies were kept moist with mineral oil, and the recording setup was enclosed in a grounded Faraday cage to minimize electrical noise. All stimulation experiments were conducted under anesthesia.

**Figure 1:**
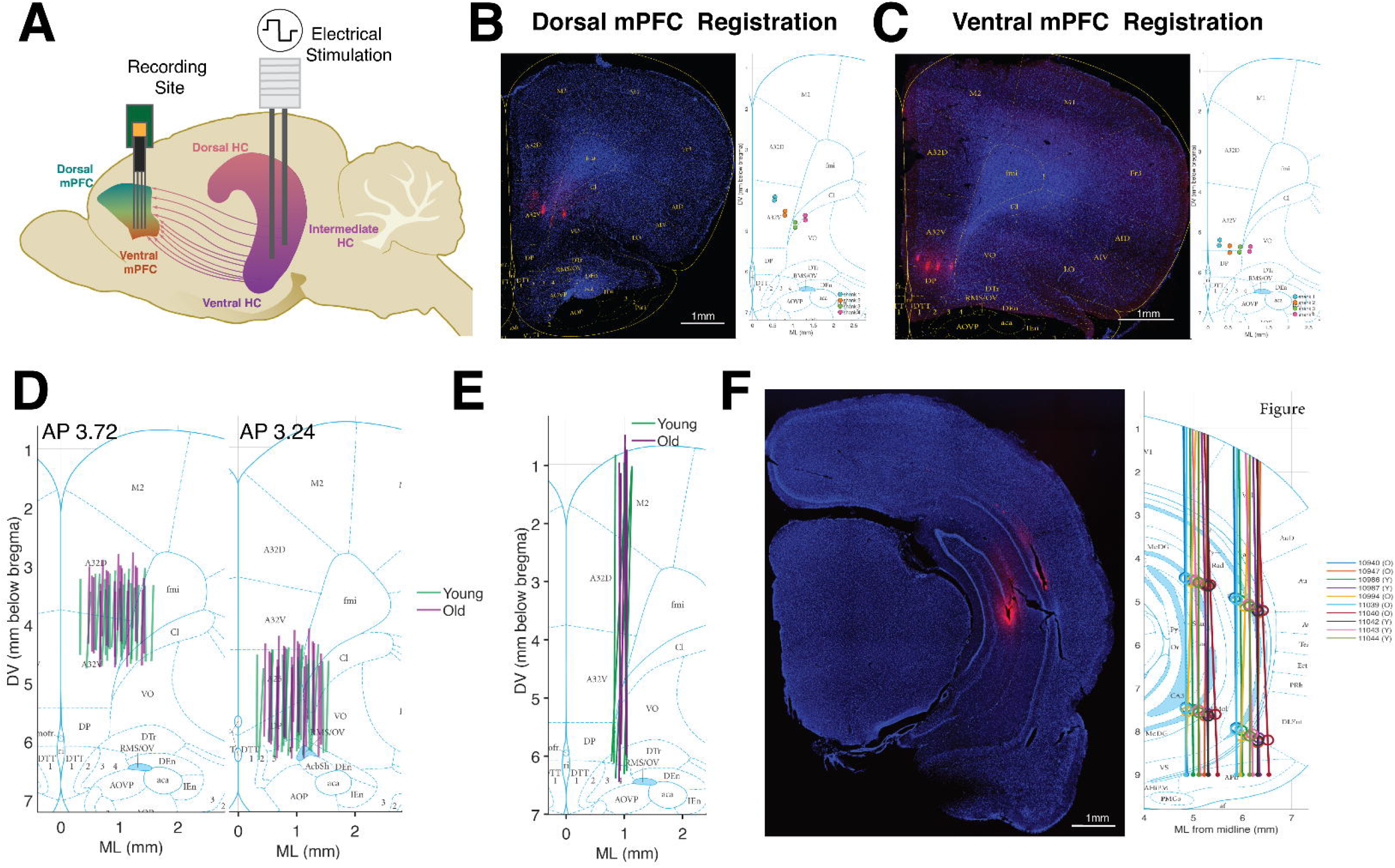
Histological Verification of probe coordinates in mPFC and HC. **A)** Schematic of stimulation and recording sites. Electrical stimulation was delivered to the intermediate (iHC) and ventral hippocampus (vHC). Neural responses were recorded across the dorsal and ventral mPFC regions reflecting direct monosynaptic input from iHC and vHC. **B)** Example of a coronal section of mPFC (*left*) stained with DAPI in blue. Fluorescent red (DiI) indicates the location of the tip of the Neuropixels 2.0 probe (4 shanks). The section is overlaid with appropriate coronal plate in yellow (Paxinos and Watson, 2017) to align the histology using anatomical landmarks. Once aligned, the probe locations are marked on the coronal plate (*right*). Here the registered probe locations correspond to the dorsal mPFC (dmPFC). **C)** Example of registration of probe location to ventral mPFC (vmPFC) **D)** Reconstructed probe tracts across all rats are projected onto the mean coronal plate for dmPFC (AP =+3.72 mm; *left*) and vmPFC (AP= +3.24 mm; *right)* respectively. In all rats, atleast 3 shanks traversed the dmPFC including A32d/A32v (prelimbic) into vmPFC including A25 (infralimbic)/dorsal peduncular cortex (DP) based on the AP coronal slide. **E)** For the comparison of the field potential responses across rats we minimized the variance in anatomical recording location by selecting the shank closest to the median ML distance from midline (0.948 mm) for each rat. The location of the selected shanks for young (green) and old (purple) rats is shown on the mPFC section with no difference in ML distribution with age (p=0.69) **F)** An example of a histological coronal section depicting the location of the bipolar stimulation electrode (1mm separation) across the CA1 region of the iHC (*left*). The reconstructed positions of all stimulation electrodes across all rats are shown on the AP slice (*right*). The circles mark the iHC and vHC stimulation coordinates for each rat as measured from the brain surface.

### Histology and microscopy

Prior to insertion, both stimulating and recording electrodes were coated with DiI (DiIC18(3), Invitrogen, Thermo Fisher Scientific) for fluorescent visualization across mPFC layers and regions. Following the experiment, rats were deeply anesthetized and transcardially perfused to extract their brains which were fixed in 4% paraformaldehyde for 24 hours, then cryoprotected in 30% sucrose for at least 3 days. From hemisected brains, coronal sections were cut on a cryostat at 40 µm and counterstained with DAPI (Vectashield H-1500-10, Vector Laboratories) before mounting onto positively charged slides (VWR SuperFrost Plus, Avantor). All slides were imaged using a slide-scanning microscope with a 5x objective (Leica DMI6000B) and 5Mpix camera (Leica DCF450). Image processing involved adjusting the brightness and contrast using ImageJ (**Figure 1C**).

### Histological Reconstruction

To verify the anatomical location of the stimulation electrode and recording probes, the histological slices were aligned with the most appropriate coronal Paxinos slide using visual inspection (Paxinos & Watson, 2017). The MATLAB image processing toolbox was used to extract the regional trace and overlay it onto the histology. The Paxinos trace was altered in 3 dimensions to best fit the histological slice using anatomical markers such as the midline, piriform cortex layer 1, corpus callosum, rhinal fissure, and olfactory ventricle in the mPFC and the CA1/CA3 cell layer in the hippocampus (**Figure 1B - C**). The probe location was then marked on the histological slice and its position interpolated onto the Paxinos slide. For each rat, 4-8 slices in the mPFC and 2-3 slices in the HC were used with the probe location at different depths to linearly interpolate the probe position.

The ML tilt and offset for each shank of the probe was determined to calculate the estimated electrode coordinates from the brain surface. Given the large inter-slide spacing of Paxinos coronal slides (∼0.5 mm), we were unable to accurately estimate the AP tilt of the probe. There was no age difference in AP placement detected in dorsal mPFC (Y = 4.296 mm ± 0.096, O = 4.200 mm ± 0.215, Wilcoxon rank-sum p = 1.00) or ventral mPFC (Y = 3.336 mm ± 0.096, O = 3.288 mm ± 0.118, Wilcoxon rank-sum p = 1.00; **Figure 1D**). The AP separation between dorsal and ventral mPFC is likely exaggerated due to the large spacing of the Paxinos coronal slides. There was also no significant age difference in the estimated ML (Y = 0.953 mm ± 0.088, O = 0.947 mm ± 0.039, p = 0.92) or DV (Y = 5.112 mm ± 0.058, O = 5.298 mm ± 0.091, p = 0.31) probe coordinates.

To restrict recordings to the mPFC, shanks located more than 1.5 mm lateral to the midline were excluded from all analyses (3 of 40 shanks). Because hippocampal projections to the mPFC differ across cortical layers, evoked field potential analyses used one shank per rat at a matched mediolateral position, sampling approximately the same layers: for each rat, the shank closest to the median ML position across all rats (0.948 mm from midline) was selected (**Figure 1E**). The ML position of the selected shank did not differ between age groups (Young: 0.936 ± 0.076 mm, Old: 1.017 ± 0.036 mm, p = 0.69). Neurons from all valid shanks were included in the single-unit analyses. Reconstructed HC stimulation electrode positions also did not differ between age groups (AP: p = 1.00, ML: p = 0.81, iHC DV: p = 0.74, vHC DV: p = 0.54; **Figure 1F**).

There were notable differences between anatomical atlases with respect to the coordinates of mPFC subregions. In Paxinos and Watson (7th ed.), the prelimbic region or A32v spans 3.6 to 4.2 mm from the brain surface at AP 3.00 to 4.68 mm, differing from earlier editions of the atlas. The region ventral to A32v (prelimbic) is labeled as either A25 (infralimbic cortex) or DP (dorsal peduncular cortex) depending on the coronal section, whereas Swanson’s atlas (Swanson, 2018), does not delineate the DP region. The AP transition from the medial orbital cortex to A25/DP also varies between the atlases (Paxinos and Watson: AP: 3.72 mm, Swanson: AP: 3.9 mm). Given this variability, we did not assign electrodes to individual subregions. Instead, following the dorsal and ventral mPFC groupings used in prior work (Gabbott et al., 2005; Laubach et al., 2018; Narayanan & Laubach, 2009; Diehl & Redish, 2023) electrodes between 2.1 and 3.8 mm from the brain surface were classified as dorsal mPFC (dmPFC, corresponding to A32/prelimbic) and electrodes deeper than 3.8 mm as ventral mPFC (vmPFC, corresponding to A25/DP). The 3.8 mm boundary was the mean dorsoventral separation between these regions across the coronal sections spanned by our recordings.

### Data Acquisition

Data collection began at least 20 minutes after probe insertion to allow for tissue stabilization. Signals were recorded using the Neuropixels acquisition system (Jun et al., 2017) and SpikeGLX software. Signals from the Neuropixels probe were amplified at a gain of 80, band-pass filtered between 0.5 Hz and 10 kHz and digitized at 30 kHz. The signal was then multiplexed and transmitted to a Neuropixels PXIe acquisition system via a 5 m tether cable. After recording a 10-minute baseline, 25 biphasic pulses (100 - 600 µA, randomized amplitude, 0.5 ms pulse width) were delivered to CA1 of iHC using a Digitimer DS4 Bi-phasic Stimulus Isolator (Digitimer Ltd.) with each pulse separated by 30-second intervals. Stimulation of the iHC was followed by a 10 min break, following which the stimulation electrode was lowered to the vHC. The same sequence of stimulation pulses delivered to the iHC were then delivered to the vHC.

### Spike Sorting

Electrical stimulation artifacts (0 - 2 ms post-stimulus) and mechanical artifacts from stimulation electrode repositioning (during lowering from iHC to vHC) were removed from the raw data prior to spike sorting using custom MATLAB scripts and imported functions (https://github.com/djoshea/neuropixel-utils). The excised timestamps were later realigned after spike sorting (marked with red arrow in **Figure 2A**).

**Figure 2:**
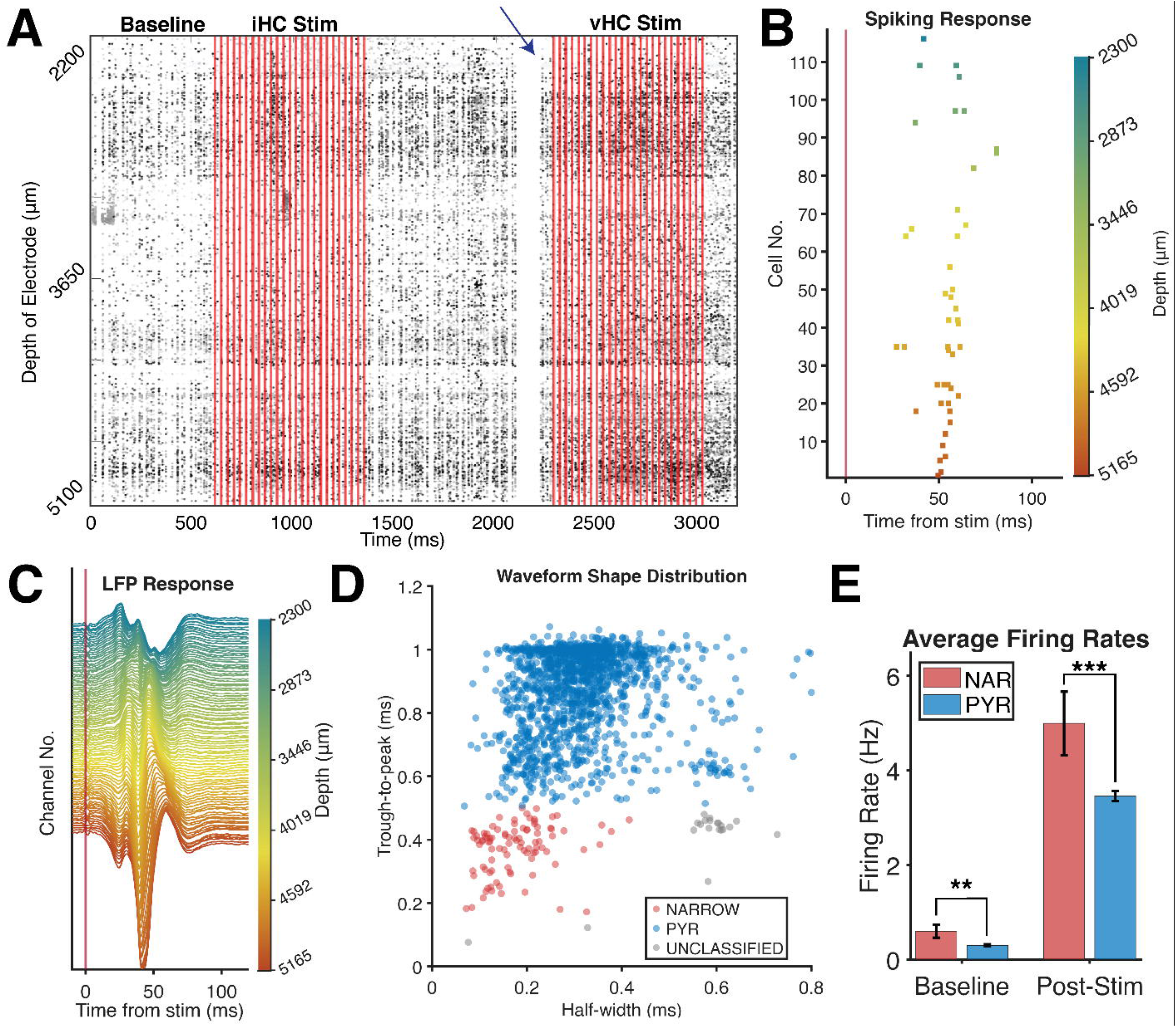
Evoked Local Field Potential and Single Unit mPFC responses to stimulation. A) Spiking activity of mPFC neurons (n=207) on each row organized by depth across the experiment in one rat. The vertical red lines represent 25 stimulation pulses (100-600 µA; 30 s apart) delivered to iHC in a randomized order. After (10 min) the stimulation probe was lowered from iHC to vHC coordinates and the pulses were delivered in the same order. To remove any artifacts from lowering the stimulation probe during recording, this epoch was removed prior to spike sorting (blank section indicated by arrow). **B)** Response of mPFC neurons across the dorsoventral gradient (blue = dorsal, orange = ventral) to a single stimulation pulse in a young rat. Coordinated neural response across the mPFC is greatest ∼50ms post stimulation. **C)** The response of evoked local field potentials (fEPSP) to the same stimulation pulse with depth indicated by the same color gradient. Here, the fEPSPs are stronger in ventral regions than dorsal. **D)** Waveform-based classification of neurons (n=1661) into narrow waveform neurons (n=94, putative interneurons; *red*) and putative pyramidal cells (n=1454, PYR; *blue*) and unclassified (n=113,*grey*). Individual waveform traces shown in red and blue, respectively. Only wide-waveform PYR neurons are included in analysis as interneuron subtypes cannot be determined. **E)** The average firing rate of the narrow waveform and pyramidal neurons in the baseline and post-stim period reveal a lower firing rate of the wide-waveform putative pyramidal cells compared to the narrow waveform neurons. **p<0.01,***p<0.001.

The automated spike-sorting algorithm Kilosort4 (Pachitariu et al., 2024) was used to identify an initial set of clusters (putative neurons) followed by manual curation using Phy (https://github.com/cortex-lab/phy). Clusters were excluded based on the following spike-sorting quality metrics: mean waveform amplitude < 35 µV, > 2% of spikes falling within inter-spike intervals of 2 ms, n < 100 spikes, maximum drift >30 µm, or if baseline firing rates were not between 0.01 - 20 Hz.

### Cell Type Classification

Units were classified as putative pyramidal or narrow-waveform neurons based on extracellular waveform characteristics (Barthó et al., 2004). Waveforms were extracted from the channel with maximum amplitude and up-sampled from 30 kHz to 250 kHz using interpolation to increase temporal resolution for spike-width measurement. Trough-to-peak duration (time interval from the peak negative deflection to the subsequent positive peak) and half-width (duration at 50% of trough amplitude) were then measured. Six percent of total units were classified as narrow-waveform neurons (NAR; n = 94) if trough-to-peak duration was 0.16 - 0.59 ms and half-width < 0.4 ms, and 87% of units with trough-to-peak duration 0.60 - 1.13 ms were classified as putative pyramidal neurons (PYR; n = 1454; **Figure 2D**). These threshold ranges are based on established values for rodent medial prefrontal cortex (Barthó et al., 2004) and can distinguish between fast-spiking narrow-waveform neurons from regular-spiking excitatory neurons. Seven percent of units not meeting criteria for either category were labeled as unclassified (n = 113). Only putative excitatory neurons were present in sufficient numbers across rats to be included in subsequent single-unit analyses. Consequently, the responses of NAR and unclassified neurons to stimulation were not analyzed given the low sample size. Measured baseline firing rates were low in the present study, which was likely due to anesthesia. Even so, firing rates of the NAR neurons (putative interneurons) were higher than PYR during post-stimulation periods (5 - 100 ms: NAR: 4.99 ± 0.67 Hz vs PYR: 3.46 ± 0.11 Hz; p = 0.001; **Figure 2E**) suggesting that putative interneurons maintained higher spontaneous activity compared to pyramidal cells, consistent with physiological properties of these neurons (Barthó et al., 2004).

### LFP Analysis

The initial 10 min recording period was used to detect anomalous channels (noisy or dead) as described in International Brain Laboratory et al., (2022) which were excluded from subsequent analysis. Briefly, noisy channels were identified as having an average power-spectral density in the 12 - 15 kHz range (0.8 x Nyquist) > 0.02 µV^2^Hz^-1^ or more than twice the median value. A total of 3 channels across all rats were identified as noisy and eliminated from analysis. Low-amplitude (dead) channels were detected by computing the zero-lag cross-correlation peak with the common median reference. Subtracting a channel’s median trend from this cross-correlation should be ∼0 for normal channels and channels with a value less than -0.5 are considered dead. A total of 2 channels out of 384 per rat were excluded as dead channels.

For each channel on the selected LFP shank (*see Histological Reconstruction*), the mean voltage of the local field potential (LFP) during 2 min of pre-stimulation baseline was subtracted to remove the DC offset. The LFPs then underwent common-average referencing across all channels to remove 60 Hz noise and electrical artifacts. The LFP was down-sampled to 2 kHz and low-pass filtered (500 Hz) with a 12th-order Butterworth filter. To reduce stimulus artifact, data spanning 0.3 ms before to 0.1 ms after each pulse was removed. A total of 13 of 500 (2.6%) stimulation pulses (5 in young, 8 in old) were not analyzed due to observed electrical noise artifacts pre-stimulation. These trials were also excluded from single-unit analyses.

### Stimulation Response Curves

To characterize the relationship between stimulus intensity and magnitude of evoked response, we examine the amplitude of the initial (monosynaptic) negative deflections in the LFP representing the excitatory field potentials (fEPSP). For each rat, we determined the peak amplitude per session by averaging across the 3 channels with the lowest peak amplitude in the monosynaptic window (5-35 ms), to ensure there was no outlier channel. We fit a least-squares logistic function (MATLAB, Optimization Toolbox function: *lsqcurvefit*) to the amplitude-intensity relationship and determined the stimulation current at which the fEPSP response magnitude reached 80% of maximum response (i80; **Figure 3C**). This i80 metric helps account for differences in the fEPSP response amplitude due to variability in stimulation electrode placement between rats (Takita et al., 1999, 2010). Stimulus-response curves were generated separately for iHC and vHC stimulation trials for each rat and fEPSP amplitude showed an increase with current intensity across all rats (**Figure 3D**) .

**Figure 3.**
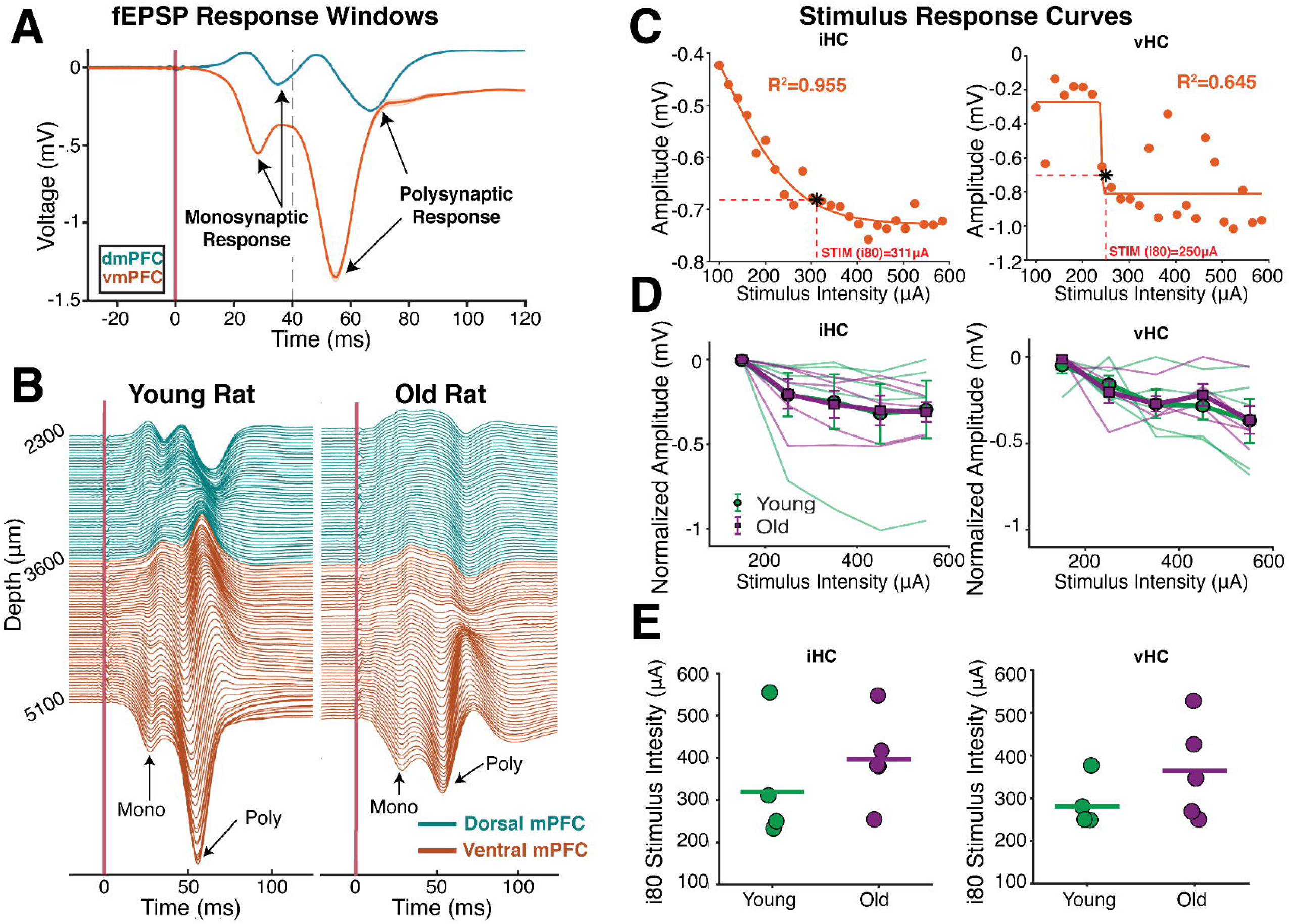
mPFC monosynaptic fEPSP response amplitude increases with stimulus intensity. **A)** Illustrative example from a single young rat of the short-latency monosynaptic and longer latency polysynaptic evoked field potentials (fEPSPs) from dorsal (dmPFC) and ventral (vmPFC) channels in response to a single stimulation. Arrows indicate the peak fEPSP (negative deflection) in the monosynaptic (5-35ms) and polysynaptic (35-100ms) window. **B)** vmPFC (*orange*) and dmPFC (*blue*) fEPSP traces averaged across all stimulation intensities for one young and one old rat. Both age groups show two distinct peaks in vmPFC corresponding to mono- and poly-synaptic population responses as denoted by arrows. **C)** Example stimulus-response curves from a young rat showing monosynaptic peak amplitude increases with stimulus intensity in both iHC (left) and vHC (right) regions. Response amplitude was fit to a logistic function, and stimulus intensity at 80% of maximum response (i80) was determined (red dashed line). **D)** Stimulus-response curves of mPFC monosynaptic amplitude for young (green) and old (purple) rats normalized to initial amplitude. All rats show an increase in fEPSP amplitude with increased stimulation intensity, though the magnitude of this increase varies across individuals. Mean ± SEM shown. **E)** To account for differences in stimulation location between rats, the i80 stimulation amplitude was determined for each rat. There was no difference in the stimulation intensities to reach plateau responses (i80) between age groups (iHC: t(8) = 0.44, p = 0.67, vHC: t(8) = −0.49, p = 0.64).

For all comparative analyses, we selected 5 consecutive iHC and vHC stimulation trials beginning at the i80 intensity for each rat (**Figure 3E**). To account for fEPSP peaks occurring at different depths across the recording region, we constructed a template for each mPFC region, layer, and HC stimulation site by identifying the largest magnitude (most negative) response at each timepoint across all channels within a 5 - 100 ms post-stimulus window, averaged across the 5 i80-intensity trials. Visual examination of these templates revealed that monosynaptic peaks occurred between 20 - 30 ms, while polysynaptic peaks ranged from 38-80 ms. Using these more confined latency windows, we identified the channel with the maximum (lowest) peak amplitude per condition (averaged across the 5 i80-intensity trials). We then selected all channels in that region and condition with peak amplitudes ≥ 50% of the maximum amplitude. We averaged responses across selected channels to determine final amplitude and time-to-peak values for statistical comparisons.

### Neuron Analysis

For each neuron, stimulus-evoked responses were quantified as baseline-subtracted firing rate that was calculated as the difference between response window firing rate and pre-stimulus baseline (200 to 0 ms), averaged across all trials within each stimulation site. For comparative analysis, we restricted analysis to the 5 stimulation trials following i80 stimulation intensity. Change in firing rate was calculated in two post-stimulation time windows: monosynaptic (5 - 35 ms) and polysynaptic (35 - 100 ms) corresponding to the rise and fall of the fEPSPs and verified by examination of the peristimulus time histograms (PSTH; **Figure 4**). We also calculated the percentage of neurons that were recruited in the post-stimulation window (0-100ms) with an evoked firing rate ≥ 1 Hz relative to baseline.

**Figure 4.**
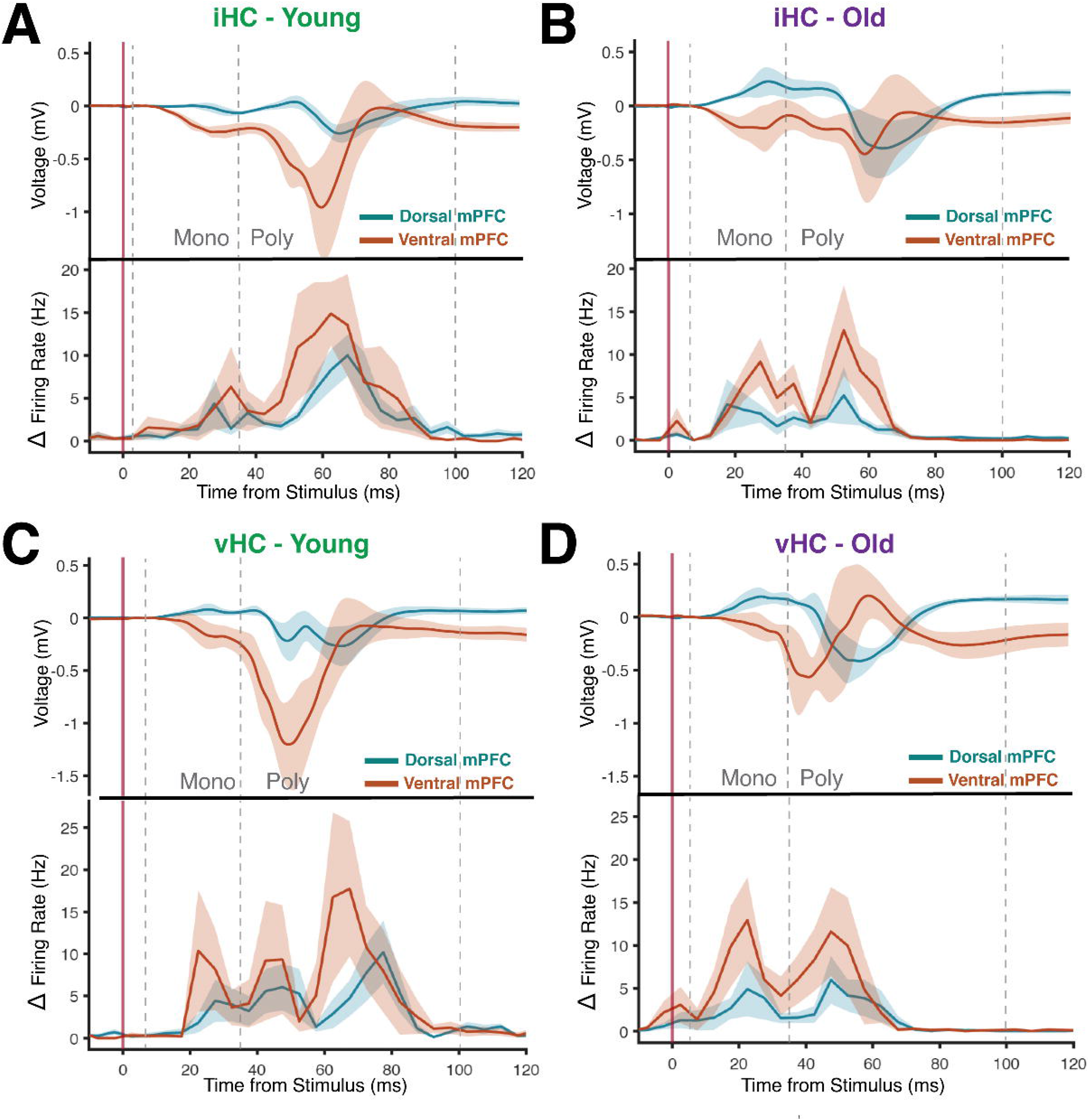
Stimulation evoked fEPSP and single-unit responses show distinct monosynaptic and polysynaptic responses. The average evoked amplitude of fEPSP response (*top*) and evoked firing rate (*bottom*) of dorsal (*blue*) and ventral (*orange)* mPFC regions to stimulation in young (n=5) and old (n=5) rats. **A**) iHC stimulation in young rats. An initial response peak in ventral mPFC fEPSP amplitude and evoked firing is seen in the monosynaptic (5-35ms) window (*dashed vertical line)*, followed by a larger response peak in the polysynaptic (35-100ms) window. The response in dorsal regions is lower than ventral in both monosynaptic and polysynaptic windows. **B)** iHC stimulation in old rats. Similar to the young rats, the average evoked responses show a bimodal distribution in the mono- and poly-synaptic windows and is greater in the ventral compared to dorsal regions. Unlike the young rats, the polysynaptic ventral mPFC responses are not greater than the monosynaptic response. **C)** vHC stimulation in young rats. Average evoked responses in dorsal and ventral mPFC are similar to iHC stimulation in both monosynaptic and polysynaptic windows. The timing of averaged evoked firing response peaks are more distributed with vHC stimulation compared to iHC stimulation, particularly in the polysynaptic window where 2 distinct peaks emerge. **D)** vHC stimulation in old rats. The average fEPSP and evoked firing response to vHC stimulation in mono- and polysynaptic windows are similar to iHC stimulated responses in dorsal and ventral mPFC.

### Statistical Analysis

All analyses treated the rat as the unit of analysis (5 young, 5 old). Channel-level and unit-level measurements were first averaged into one value per rat per session per condition. Linear mixed-effects models were used to test the effects of age, HC stimulation site, mPFC region, and response window (fixed effects) on LFP response amplitude, LFP time-to-peak, evoked firing rate, and the percentage of recruited neurons, with rat as a random intercept. Models were fitted by restricted maximum likelihood using the MATLAB function *fitlme*. For mPFC depth analysis, channels and units were averaged into 100 µm depth bins and entered as a continuous predictor to the model. Planned contrasts were specified a priori based on two grounds. First, distinct iHC and vHC projections to the dmPFC and vmPFC (Hoover & Vertes, 2007; Alemán-Andrade et al., 2025). Second, distinct circuits are involved in the monosynaptic and polysynaptic responses, namely the direct HC input and local mPFC recruitment, respectively (Jay et al., 1992; Taylor et al., 2016). Each combination of stimulation site and response window was therefore treated as a distinct functional pathway that could be differentially altered by age. Contrasts comparing: *Monosynaptic vs polysynaptic* responses for iHC and vHC per age group, *Dorsal vs ventral mPFC* responses for iHC and vHC per age group, *Young vs Old responses* per mPFC region, window and stimulation site, were tested on model coefficients (*coefTest*) and corrected using the Holm-Bonferroni procedure. All statistical tests used a significance threshold of α = 0.05. Effect sizes are reported as η² for model terms and as partial η² (η²p) for planned contrasts. All analyses were performed in MATLAB R2025b (The MathWorks, Natick, MA), and data are presented as mean ± SEM unless otherwise noted.

### Code Accessibility

All custom scripts used here for analysis can be found at https://github.com/Barnes-Lab/HC_PFC_Stim_Neuropixel which is publicly available.

## Results

### mPFC monosynaptic fEPSP response amplitude increases with stimulus intensity

Electrical stimulation delivered to the CA1 region of intermediate (iHC) and ventral hippocampus (vHC) evoked LFP and single-unit responses in the mPFC (**Figure 2**). Negative deflections, thought to reflect extracellular excitatory field potentials (fEPSPs), were observed along the mPFC dorsoventral gradient and were strongest in the ventral region **(Figure 3A-B).** We also observed a stimulus-evoked positive deflection in the LFP that may represent either an inhibitory response or a passive return current along the local dendritic projections (Leung, 2011; Buzsáki et al., 2012). As the recording locations were parallel to the local dendritic projections, we cannot draw any direct inferences about dendritic sinks and sources and thus do not consider them in this analysis.

We observed two distinct temporal components of the fEPSP across channels: an initial response, thought to be generated by the monosynaptic iHC/vHC projection and a secondary longer latency peak, thought to be a polysynaptic response generated by local mPFC feed-forward circuits (**Figure 3A-B**). With increasing stimulation intensity (100 - 600 µA), the peak amplitude of the monosynaptic fEPSP response increased up to a plateau in both young and old rats, indicating a saturation of monosynaptic responses (**Figure 3C-D**). We fit a least-squares logistic function to the amplitude-intensity relationship for each rat and determined the stimulation current at which the fEPSP response magnitude reached 80% of maximum response (example of logistic fit and i80 stimulation amplitude calculation for iHC and vHC in **Figure 3C**). There was no difference in the stimulation intensities to reach 80% of maximum response (i80) between age groups (**Figure 3E**; iHC: t(8) = 0.44, p = 0.67, vHC: t(8) = −0.49, p = 0.64). Subsequent comparisons of fEPSP and single-unit responses averaged responses from 5 consecutive stimulation trials, starting at the i80 stimulation amplitude for each rat per condition.

### HC stimulation evokes temporally distinct monosynaptic and polysynaptic mPFC responses

Both iHC and vHC stimulation produce temporally distinct mPFC fEPSP response peaks corresponding to distinct synaptic components: a monosynaptic peak (time-to-peak between 5 - 35 ms) and polysynaptic peak (time-to-peak between 35 - 100 ms; **Figure 4**), with latencies on par with previous studies (Laroche et al., 1990; Takita et al., 2010; Taylor et al., 2016). Stimulation-evoked single-unit firing also showed similar monosynaptic and polysynaptic latencies (**Figure 4**).

In young rats, iHC stimulation produced an initial smaller monosynaptic fEPSP and evoked firing response (peak ∼28 - 30 ms) followed by a larger polysynaptic response (∼60 ms). The response in ventral regions was greater than dorsal regions in both mono- and poly-synaptic windows (**Figure 4A**). In old rats, the ventral mPFC response to iHC stimulation was greater than dorsal mPFC, but there was no difference in the magnitude of monosynaptic (peak ∼25 - 27 ms) and polysynaptic (∼55 - 60 ms) responses (**Figure 4B**). Average vHC stimulation-evoked fEPSP responses in young rats were similar to iHC stimulation with smaller monosynaptic (peak ∼22 ms) and larger polysynaptic (peak ∼52 ms) response amplitude. While the monosynaptic evoked firing was aligned to the fEPSP peak, the average polysynaptic evoked firing occurs in two distinct peaks (∼45 ms and 65 ms) (**Figure 4C**). In old rats the average fEPSP and evoked firing response to vHC stimulation in monosynaptic (peak ∼20-22 ms) and polysynaptic (peak ∼42-50 ms) windows were similar to iHC stimulated responses in dorsal and ventral mPFC.

It should be noted that one study reported a time-to-peak of the initial fEPSP of ∼20 ms (Takita et al., 1999) while we observed latencies between 22 and 30 ms. This difference might be explained by the fact that prior studies used single-electrode measurements. Our analysis measured the average peak fEPSP response across multiple recording sites thus reflecting the maximal population response rather than response onset or maximal slope and was chosen due to the high variability in the onset and slope across recording depths. Synaptic conduction timing may also be affected by the exact placement of the stimulating electrodes, differences in the type of anesthesia, or body temperature between studies (Volgushev et al., 2000).

### Evoked mPFC polysynaptic responses are greater than monosynaptic in young but not old rats

Since the monosynaptic response reflects direct HC input (Jay et al., 1992; Laroche et al., 1990), and the polysynaptic response reflects recruitment of local mPFC circuits (Dégenètais et al., 2003; Izaki et al., 2003; Taylor et al., 2016), we predicted that the two components may be differently affected by age. fEPSP amplitude in each window was therefore modeled with a linear mixed-effects model with age, HC stimulation site, mPFC region, and response window as fixed effects and rat as a random intercept (**Table 1**). The polysynaptic fEPSP amplitude was larger than the monosynaptic amplitude (**Figure 5A**, **Table 1**) qualified by a significant Age x mPFC region x Window interaction (**Table 1**). Because iHC and vHC projections to the mPFC are anatomically distinct, pre-specified contrasts were performed between the two windows within each site and age group. The polysynaptic increase was present in young rats at both sites (iHC: β = −1.31, Holm p = 0.008, η²p = 0.67; vHC: β = −1.44, Holm p = 0.01, η²p = 0.64) while in old rats it was smaller and did not reach significance at either site (vHC: β = −0.83, Holm p = 0.06; iHC: p = 0.38). There was no overall effect of age or stimulation site on fEPSP amplitude (**Table 1**).

**Figure 5.**
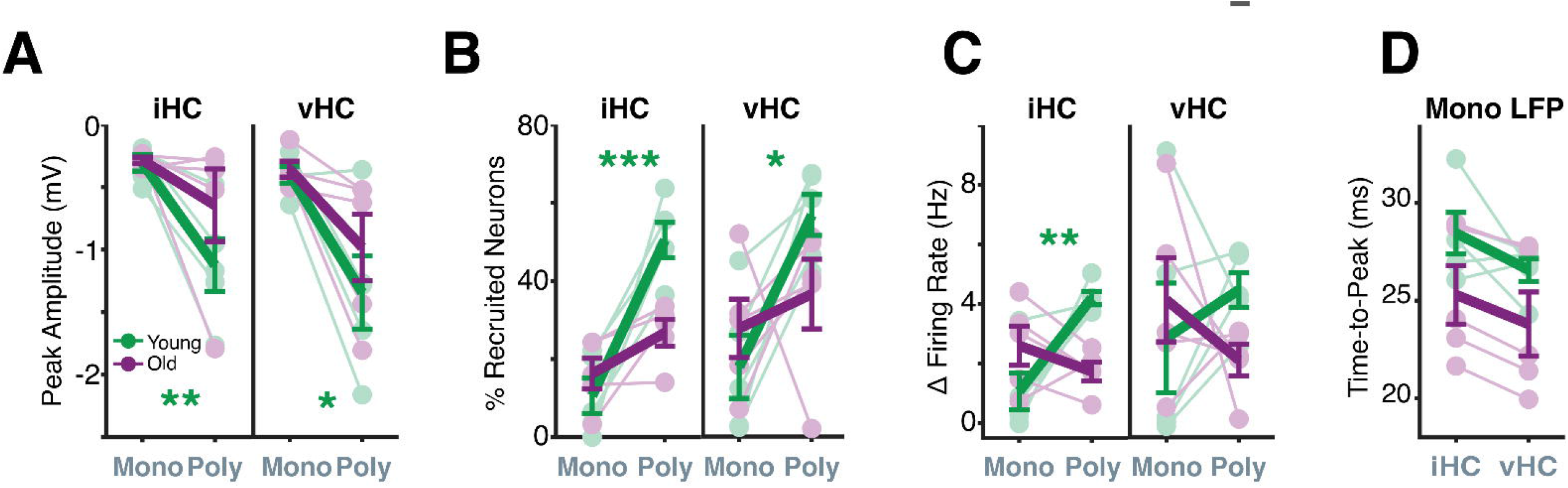
Evoked mPFC polysynaptic responses are greater than monosynaptic in young but not old rats. **A)** The peak amplitude of fEPSP responses averaged across all mPFC recording depths is greater in the polysynaptic window than in the monosynaptic window with both iHC and vHC stimulation in young rats (green, N = 5; iHC: β = −1.31, Holm p = 0.008; vHC: β = −1.44, Holm p = 0.010). In old rats (purple, N = 5), this increase did not reach significance for either site (vHC: β = −0.83, Holm p = 0.06; iHC: p = 0.38). Individual rats are shown with connecting lines; group means ± SEM are shown with thick lines and error bars. **B**) Neurons with evoked firing rates ≥ 1 Hz above baseline in the 0 - 100 ms post-stimulus window were classified as recruited (Y: n = 520, O: n = 920 neurons). In young rats, more mPFC neurons are recruited in the polysynaptic window than in the monosynaptic window with both iHC and vHC stimulation (iHC: β = 37.9, Holm p < 0.001; vHC: β = 38.0, Holm p = 0.028). This pattern is absent in old rats (iHC: Holm p = 0.065; vHC: Holm p = 0.50). C) With iHC stimulation, evoked firing increased from the monosynaptic to the polysynaptic window in young but not old rats (Y: β = 2.82, Holm p = 0.009; O: p = 0.45). With vHC stimulation, this difference was not significant in either age group (Y: Holm p = 0.54; O: Holm p = 0.54). D) The time-to-peak of the monosynaptic fEPSP did not differ between stimulation sites (Y: Holm p = 0.23; O: p = 0.61) or between age groups (iHC: Holm p = 0.29; vHC: p = 0.34). Significance indicated as *p < 0.05, **p < 0.01, ***p < 0.001. Within-age and between-age planned contrasts report Holm-corrected p values. *See* **Table 1** for the full model and interaction terms.

**Table 1:** Effects of age, HC stimulation site, mPFC region and response window on evoked responses. . Separate mixed-effects models for fEPSP amplitude, percentage of recruited PYR neurons, and evoked firing rate, each with age, HC stimulation site (vHC - iHC), mPFC region (ventral - dorsal; boundary at 3800 µm) and response window (polysynaptic - monosynaptic) as fixed effects and rat as a random intercept (n = 5 young, 5 old). β is the estimated difference for each term, in the measure’s units and effect size is η². Planned contrasts within age, site and window are reported in the text. Here fEPSP amplitude is negative thus more negative values indicate larger responses.

| Table 1. Effect of Age, HC Stimulation Site, mPFC Region and Response Window |  |  |  |  |  |  |  |  |  |
| --- | --- | --- | --- | --- | --- | --- | --- | --- | --- |
| | LFP amplitude (mV) | | | % recruited PYR | | | PYR ΔFR (Hz) | | |
| | $\beta$ | $p$ | $\eta^2$ | $\beta$ | $p$ | $\eta^2$ | $\beta$ | $p$ | $\eta^2$ |
| Age | +0.227 | 0.260 | 0.024 | -7.630 | 0.067 | 0.030 | -0.525 | 0.490 | 0.010 |
| HC Site (vHC - iHC) | -0.183 | 0.077 | 0.016 | +7.870 | 0.067 | 0.032 | +0.793 | 0.061 | 0.024 |
| mPFC Region (ventral - dorsal) | -0.651 | **0.002** | 0.200 | +12.00 | **<0.001** | 0.073 | +1.800 | **<0.001** | 0.122 |
| Window (poly - mono) | -0.687 | **0.004** | 0.222 | +25.60 | **0.001** | 0.334 | +0.758 | 0.330 | 0.022 |
| Age × HC Site | -0.047 | 0.800 | 0.000 | +4.510 | 0.560 | 0.003 | +0.270 | 0.720 | 0.001 |
| Age × mPFC Region | +0.520 | 0.120 | 0.032 | +1.580 | 0.680 | 0.000 | -0.341 | 0.630 | 0.001 |
| Age × Window | +0.388 | 0.280 | 0.018 | -32.200 | **0.015** | 0.132 | -4.310 | **0.019** | 0.174 |
| HC Site × mPFC Region | +0.078 | 0.440 | 0.001 | +0.694 | 0.850 | 0.000 | +0.932 | 0.066 | 0.008 |
| HC Site × Window | -0.194 | 0.160 | 0.004 | -0.809 | 0.920 | 0.000 | -1.040 | 0.290 | 0.010 |
| mPFC Region × Window | -0.588 | **0.015** | 0.041 | +7.090 | 0.370 | 0.006 | +0.583 | 0.600 | 0.003 |
| Age × HC Site × mPFC Region | +0.034 | 0.860 | 0.000 | +8.470 | 0.280 | 0.002 | +0.793 | 0.390 | 0.001 |
| Age × HC Site × Window | -0.141 | 0.590 | 0.001 | -2.580 | 0.870 | 0.000 | -0.320 | 0.870 | 0.000 |
| Age × mPFC Region × Window | +0.921 | **0.043** | 0.025 | -14.40 | 0.360 | 0.007 | -2.990 | 0.200 | 0.021 |
| HC Site × mPFC Region × Window | -0.211 | 0.260 | 0.001 | -1.240 | 0.890 | 0.000 | -0.966 | 0.510 | 0.002 |
| Age × HC Site × mPFC Region × Window | -0.288 | 0.430 | 0.001 | -7.380 | 0.670 | 0.000 | -2.540 | 0.390 | 0.004 |

To examine spiking output within the time windows corresponding to the monosynaptic and polysynaptic fEPSP peaks, the percentage of recruited neurons and the evoked firing rate were measured in each window and modeled with the same LME structure (**Table 1**). More neurons were recruited in the polysynaptic than the monosynaptic window in young but not old rats (**Figure 5B**, **Table 1**). Using the same pre-specified contrasts as for fEPSP amplitude within each stimulation site and age group, increased polysynaptic recruitment was observed in young rats with both sites (iHC: β = 37.9, Holm p < 0.001, η²p = 0.85; vHC: β = 38.0, Holm p = 0.03, η²p = 0.55), whereas no increase was observed in old rats with either site (iHC: Holm p = 0.065; vHC: Holm p = 0.50). Similarly, changes in evoked firing rates between monosynaptic and polysynaptic windows differed between age groups (**Figure 5C**; **Table 1**). In young rats, firing rates increased in the polysynaptic relative to the monosynaptic window with iHC stimulation (β = 2.82, Holm p = 0.009, η²p = 0.66), while no significant increase was observed for the remaining contrasts (Y vHC: Holm p = 0.54; O iHC: p = 0.45; O vHC: p = 0.54). The time-to-peak of the fEPSP did not differ between stimulation sites (p = 0.14) or between age groups (p = 0.17; **Figure 5D**).

The pattern of an initially small fEPSP peak followed by a larger amplitude fEPSP peak as seen in young rats aligns with previous characterizations of mPFC responses to vHC stimulation (Izaki et al., 2003; Laroche et al., 1990; Takita et al., 1999; Taylor et al., 2016). The absence of an increase in polysynaptic spiking response in old rats suggests a deficit in local mPFC circuit recruitment following the initial monosynaptic HC response (Dégenètais et al., 2003; Izaki et al., 2003; Taylor et al., 2016).

### Monosynaptic, but not polysynaptic, fEPSP depth gradients in mPFC are preserved with age

Both iHC and vHC project most densely to the ventral mPFC (Hoover & Vertes, 2007; Anastasiades et al., 2018; Liu & Carter, 2018) although the dorsoventral extent of their synaptic terminals differ (Alemán-Andrade et al., 2025). We therefore expected evoked responses to vary with age along the dorsoventral mPFC axis depending on the HC stimulation site and overall have a greater response in the ventral mPFC region. To examine how the evoked responses varied along the dorsoventral gradient, peak fEPSP amplitude was measured in 100 µm depth bins along the probe. This continuous depth gradient was used as an independent variable in a linear mixed-effects model to predict fEPSP amplitude along with response window, stimulation site and age as fixed effects and rat as random intercept (**Figure 6**, **Table 2**). Overall responses in the dmPFC and vmPFC categorically defined regions, modeled in **Table 1**, were additionally compared between age groups with planned contrasts within each stimulation site and window (**Figure 6**, *inset*).

**Figure 6:**
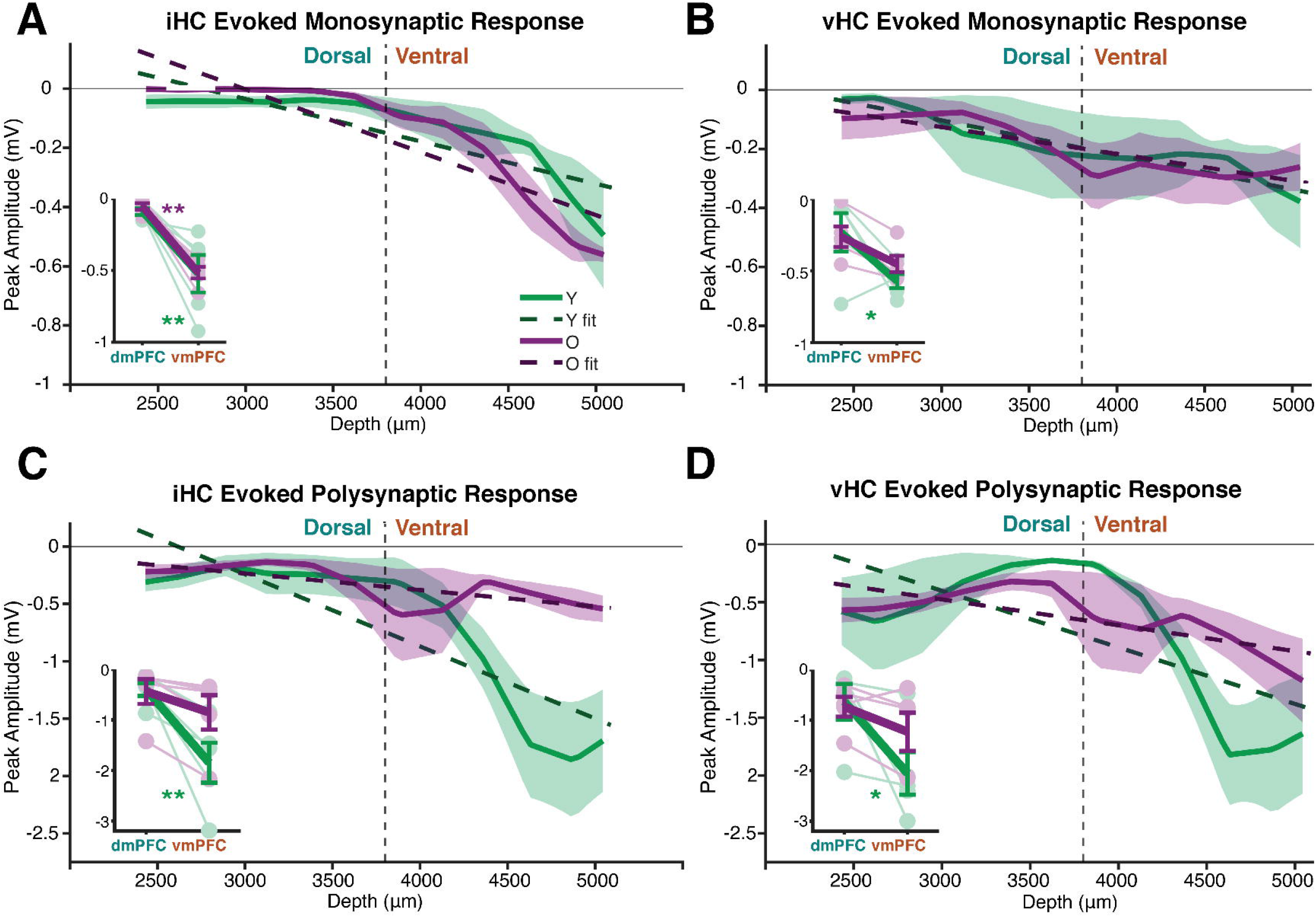
Amplitude of monosynaptic and polysynaptic peak fEPSP response across mPFC depths. Peak fEPSP response amplitude across mPFC depths (100 µm bins) for young (N = 5, *green*) and old (N = 5, *purple*) rats. Linear fits of response amplitude with depth for each age group are overlaid as dashed lines. (*Inset*) Average response in dorsal (dmPFC) and ventral (vmPFC) regions with individual rats shown with connecting lines; group means ± SEM shown with thick lines and error bars. More negative fEPSP amplitude indicates a larger response. **A)** Monosynaptic response to iHC stimulation. In both age groups, there is little response in dorsal regions with a large increase in response at ventral depths (Y: β = −0.014, Holm p = 0.002; O: β = −0.021, Holm p < 0.001). The average response amplitude in vmPFC is greater than dmPFC in young and old rats (Y: β = −0.43, Holm p = 0.003; O: β = −0.46, Holm p = 0.003), with no age difference in either region (dmPFC: p = 0.57; vmPFC: p = 0.96). **B)** Monosynaptic response to vHC stimulation. Response amplitude increases gradually with depth in both age groups (Y: β = −0.012, Holm p = 0.018; O: β = −0.009, Holm p = 0.028). The average vmPFC response amplitude is greater than dmPFC in young rats (β = −0.34, Holm p = 0.026) but not old rats (p = 0.11), with no age difference in either region (dmPFC: Holm p = 0.85; vmPFC: Holm p = 0.31). **C)** Polysynaptic response to iHC stimulation. Young rats show a steep increase in amplitude with depth (β = −0.063, Holm p = 0.002) with the greatest response at the most ventral depths, whereas old rats show no significant depth dependence (p = 0.28), with the largest responses at intermediate to ventral depths. The average vmPFC response amplitude is greater than dmPFC only in young rats (Y: β = −1.45, Holm p = 0.003; O: p = 0.22), and the depth profile differs between age groups (Age × Depth: β = 0.049, p = 0.025; Age × Region: β = 1.03, p = 0.045). **D)** Polysynaptic response to vHC stimulation. Young rats show an increase in amplitude with depth (β = −0.049, Holm p = 0.026) that is absent in old rats (p = 0.18), and both age groups also show a response at the most dorsal depths. The average vmPFC response amplitude is greater than dmPFC only in young rats (Y: β = −1.42, Holm p = 0.016; O: p = 0.26), with no difference between age groups (Age × Depth: p = 0.26; vmPFC: p = 0.37). Significance indicated as *p<0.05, **p<0.01, ***p<0.001. See *Table 2* for the full models.

**Table 2:** fEPSP amplitude across the dorsoventral axis of mPFC by stimulation site and response window. . For each stimulation site (iHC, vHC) and response window (monosynaptic, polysynaptic), peak fEPSP amplitude in 100 µm depth bins was modeled with a linear mixed-effects model with age and mPFC depth (continuous, increasing ventrally) as fixed effects and rat as a random intercept (N = 5 young, 5 old). Each row reports one model term: β in the measure’s units, with the mPFC Depth term giving the change in amplitude per 100 µm of depth, degrees of freedom, p value, and effect size (η²). The Age × mPFC Depth term tests whether the depth gradient differs between age groups. fEPSP amplitudes are negative; more negative values indicate larger responses. Per-age slopes and dorsal versus ventral contrasts are reported in the text and Figure 6.

| Table 2. fEPSP amplitude across mPFC depth |  |  |  |  |  |  |  |  |  |  |  |  |
| --- | --- | --- | --- | --- | --- | --- | --- | --- | --- | --- | --- | --- |
|  | iHC mono |  |  | vHC mono |  |  | iHC poly |  |  | vHC poly |  |  |
| | $\beta$ | p | $\eta^2$ | $\beta$ | p | $\eta^2$ | $\beta$ | p | $\eta^2$ | $\beta$ | p | $\eta^2$ |
| Age | 0.022 | 0.770 | 0.002 | 0.045 | 0.650 | 0.010 | 0.480 | 0.210 | 0.083 | 0.362 | 0.390 | 0.041 |
| mPFC Depth (per 100 μm) | -0.018 | **<0.001** | 0.890 | -0.010 | **0.003** | 0.700 | -0.039 | **0.002** | 0.710 | -0.036 | **0.011** | 0.570 |
| Age × mPFC Depth | -0.007 | 0.150 | 0.240 | 0.003 | 0.600 | 0.035 | 0.049 | **0.025** | 0.486 | 0.027 | 0.260 | 0.158 |
| Age × Site × Depth | $\beta = -0.0062$ , p = 0.44, $\eta^2 = 0.003$ | | | | | | | | | | | |
| Age × Site × Window × Depth | $\beta = -0.031$ , **p = 0.048**, $\eta^2 = 0.021$ | | | | | | | | | | | |

Consistent with our hypothesis, fEPSP amplitude was larger in ventral than dorsal mPFC (dmPFC, vmPFC; **Table 1**) which was qualified by a significant Age × mPFC region × Window interaction (**Table 1**). Along the dorsoventral axis, monosynaptic fEPSP amplitude increased with depth in both age groups with both stimulation sites (**Table 2**). With iHC stimulation, little monosynaptic response was observed at dorsal depths (dmPFC), with a large increase at the most ventral depths (vmPFC) in both age groups (**Figure 6A**), whereas with vHC stimulation smaller responses were present at dorsal depths and increased gradually along the gradient (**Figure 6B**). The larger monosynaptic fEPSP amplitude in the vmPFC aligns with our prediction based on denser hippocampal projections to this region (Hoover & Vertes, 2007; Anastasiades et al., 2018; Liu & Carter, 2018).

The depth profile of the polysynaptic fEPSP differed between age groups with iHC stimulation (**Figure 6C**, **Table 2**). Young rats showed a steep increase in response with depth, with the greatest responses at the most ventral depths (β = -0.06, Holm p = 0.002, η²p = 0.76), whereas old rats showed no monotonic depth gradient (p = 0.28), with their largest responses at intermediate to ventral depths. Accordingly, vmPFC amplitude was greater than dmPFC in young but not old rats as seen from planned contrasts (**Figure 6C**, *Inset;* Young (Y): β = -1.45, Holm p = 0.003, η²p = 0.73; Old (O): p = 0.22). With vHC stimulation, polysynaptic responses were greatest at the most ventral depths, with an additional increase at dorsal depths (**Figure 6D**, **Table 2**). This depth gradient in polysynaptic responses was significant only in young rats (Y: β = -0.05, Holm p = 0.026; O: p = 0.18), as was the increase from dmPFC to vmPFC (**Figure 6D**, *Inset;* Y: β = -1.42, Holm p = 0.016, η²p = 0.60; O: Holm p = 0.26). Thus, while the monosynaptic response increased with depth in both age groups, only young rats showed a polysynaptic difference between dorsal and ventral mPFC.

### Polysynaptic evoked firing is lower in old rats at ventral mPFC depths

To determine whether the mPFC depth gradient in fEPSP responses was reflected in spiking output, evoked firing rate was examined across depth with the same per-condition depth models. Unlike fEPSP amplitude, monosynaptic firing evoked by iHC stimulation did not change with mPFC depth in either age group (**Figure 7A**, **Table 3)** and average firing in dmPFC and vmPFC was similar (Young (Y): p = 0.95; Old (O): p = 0.95). With vHC stimulation, firing likewise did not change across the depth gradient (**Figure 7B**, **Table 3**) and average vmPFC firing did not differ from dmPFC within either age group (**Figure 7B**, *Inset;* Y: Holm p = 0.43; O: Holm p = 0.078).

**Figure 7:**
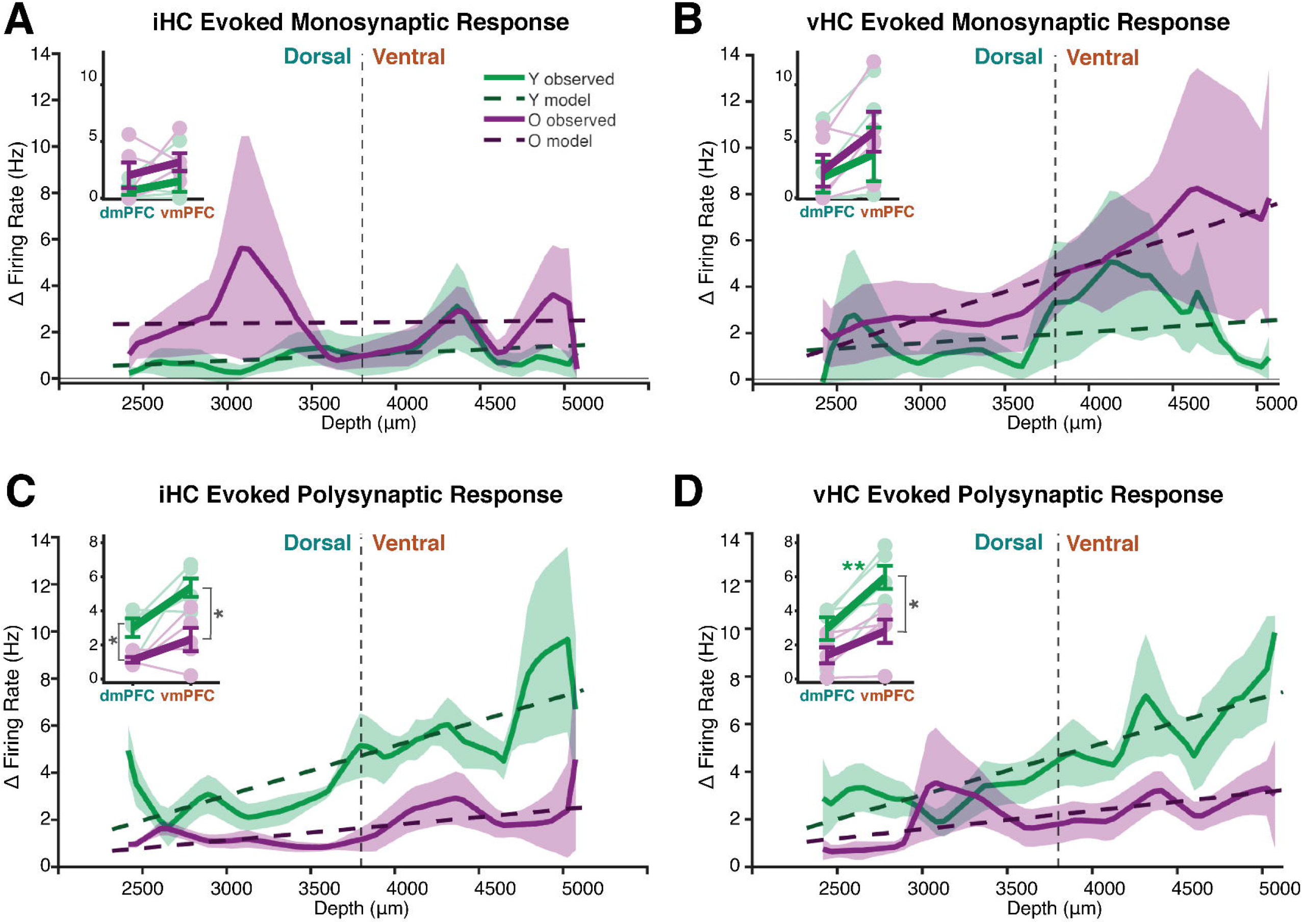
Evoked firing in monosynaptic and polysynaptic windows across mPFC depths. Average evoked firing rate relative to baseline across mPFC depths (100 µm bins) for young (N = 5, n = 520 neurons, *green*) and old (N = 5, n = 920 neurons, *purple*) rats. Linear fits of evoked firing rate with depth for each age group are overlaid as dashed lines; (*Inset*) Average response in dorsal (dmPFC) and ventral (vmPFC) regions with individual rats shown with connecting lines; group means ± SEM shown with thick lines and error bars. **A)** Monosynaptic response to iHC stimulation. Evoked firing does not change with depth in either age group (both Holm p = 1.00). Average dmPFC and vmPFC responses are similar within each age group (both Holm p = 0.95), with no age difference in either region (both Holm p > 0.57). **B)** Monosynaptic response to vHC stimulation. Evoked firing does not change significantly with depth in either age group (Y: Holm p = 0.74; O: Holm p = 0.15). Average vmPFC firing is greater than dmPFC across all rats (p = 0.047) but not within either age group alone (Y: Holm p = 0.43; O: Holm p = 0.078). **C)** Polysynaptic response to iHC stimulation. Evoked firing does not change significantly with depth in either age group (Y: β = 0.21, Holm p = 0.062; O: p = 0.48), but average vmPFC firing is greater than dmPFC across all rats (p = 0.032). Evoked firing is lower in old rats in both regions (dmPFC: β = −1.93, Holm p = 0.014; vmPFC: β = −3.53, Holm p = 0.014). **D)** Polysynaptic response to vHC stimulation. Evoked firing increases with depth in young rats (β = 0.19, Holm p = 0.003) but not old rats (p = 0.09), and average vmPFC firing is greater than dmPFC only in young rats (Y: β = 3.36, Holm p = 0.002; O: p = 0.079). The age-related reduction in evoked firing is restricted to the vmPFC (dmPFC: p = 0.085; vmPFC: β = −3.66, Holm p = 0.010). Significance indicated as *p<0.05, **p<0.01, ***p<0.001. Within-age and between-age contrasts report Holm-corrected p values. *See* **Table 3** for the full models, including interaction terms.

**Table 3:**
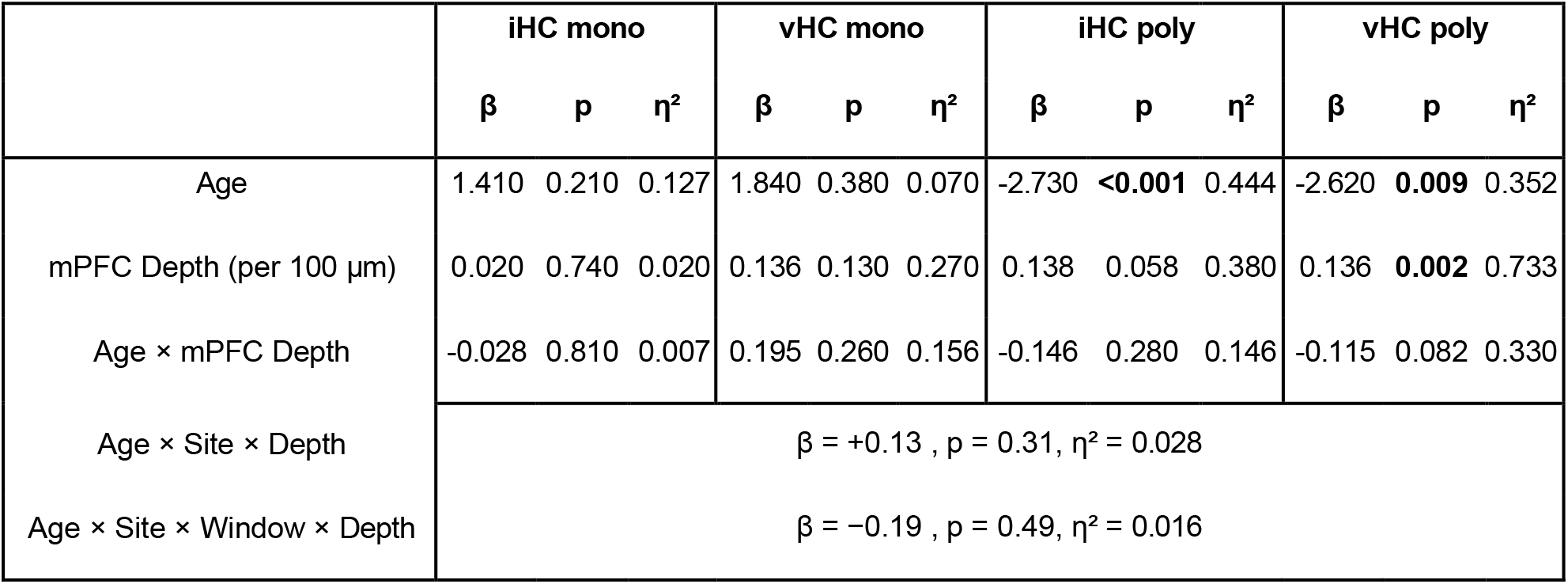
Evoked PYR firing rate across the dorsoventral axis of mPFC by stimulation site and response window. . For each stimulation site and response window, baseline-subtracted firing rate of putative pyramidal neurons was averaged into 100 µm depth bins and modeled with a linear mixed-effects model with age and mPFC depth (continuous, increasing ventrally) as fixed effects and rat as a random intercept (N = 5 young, 5 old; n = 520 young and 920 old neurons). Each row reports one model term: β, with the mPFC Depth term giving the change in firing rate (Hz) per 100 µm of depth, degrees of freedom, p value, and η². Positive slopes indicate greater evoked firing at ventral depths. Per-age slopes and dorsal versus ventral and age contrasts are reported in the text and Figure 7.

In the polysynaptic window, evoked firing was lower in old than young rats with both stimulation sites (**Table 3**). With iHC stimulation, firing did not increase significantly across the gradient (**Figure 7C**, **Table 3**), although average firing in vmPFC was greater than in dmPFC across all rats (**Figure 7C**, *Inset;* p = 0.032). With vHC stimulation, evoked firing increased steadily across the gradient (**Figure 7D**, **Table 3**), driven by young rats (Y: β = 0.19, Holm p = 0.003, η²p = 0.74; O: β = 0.08, p = 0.09), and the increase in average firing from dmPFC to vmPFC was likewise seen only in young rats (**Figure 7D**, *Inset;* Y: β = 3.36, Holm p = 0.0015, η²p = 0.78; O: β = 1.28, Holm p = 0.08). The age difference in vHC polysynaptic firing was restricted to the vmPFC (dmPFC: p = 0.09; vmPFC: β = −3.66, Holm p = 0.010, η²p = 0.65). Overall, polysynaptic firing in the vmPFC was lower in old than young rats with both stimulation sites, whereas monosynaptic firing was unaffected by age, and an increase in evoked firing from dorsal to ventral mPFC was seen only in young rats.

## Discussion

The most important findings of the present study are that, contrary to our original predictions, aging did not alter the monosynaptic functional connection between the hippocampus and mPFC, whereas the polysynaptic responses that follow this direct input were reduced in old rats. In young rats, polysynaptic fEPSP and single-unit responses were larger than the monosynaptic responses and were greater in ventral than in dorsal mPFC. Old rats showed lower evoked firing than young rats and neither the polysynaptic increase nor the ventral-dorsal difference was present. These age-related changes further differed between the iHC and vHC stimulation sites, consistent with their distinct projections to the mPFC.

### Hippocampus-evoked monosynaptic responses are unaltered with age

Surprisingly, aging did not alter the functional monosynaptic connection between the hippocampus and mPFC. We found no significant difference between young and old rats in the HC evoked monosynaptic responses in the mPFC as measured by fEPSP time-to-peak (**Figure 5D**), amplitude (**Figure 6A - B**), and evoked neural firing (**Figure 7A - B**). There was also no significant difference in the stimulus intensity at which young and old rats reached the maximum fEPSP amplitude (**Figure 3E**). An age-related reduction in the functional connectivity of this circuit was expected, given the changes observed across species in behavioral tasks presumed to engage the HC-PFC circuit (Barnes, 1987; Bizon et al., 2012; Beas et al., 2017; Kapellusch et al., 2018; Barense et al., 2002; Glisky, 2007). In primates, age-related changes in episodic and working-memory correlate with reduced volume (Shamy et al., 2011), disruption of white matter tracts (Metzler-Baddeley et al., 2011; Gunbey et al., 2014; Stadlbauer et al., 2008), and reduced functional connectivity (Ankudowich et al., 2019; Blum et al., 2014; Damoiseaux et al., 2016; Grady et al., 2012; Vidal-Piñeiro et al., 2014) between the HC and PFC. While no studies that we are aware of have examined the effect of age on this circuit in rodents, the functional connectivity of HC (Barnes, 1994; Barnes & McNaughton, 1980; Trompoukis et al., 2022) and mPFC (Allard et al., 2012; Haberman et al., 2019) projections to other brain regions have been shown to be altered with age. The current findings indicate that the monosynaptic functional connection of the HC-mPFC circuit is preserved with age, however, we cannot directly assess the structural integrity of this circuit in aged rodents.

### Ventral mPFC polysynaptic responses evoked by HC stimulation are reduced in old rats

Polysynaptic responses arise from local collaterals of mPFC neurons that receive HC input (Jay et al., 1992; Little & Carter, 2012; Anastasiades & Carter, 2021; Taylor et al., 2016). Evoked pyramidal neuron firing in the polysynaptic window was lower in old than in young rats with both iHC and vHC stimulation, particularly in vmPFC regions (**Figure 7C - D**). Consistent with this reduction, the polysynaptic fEPSP and single-unit responses that were greater than the monosynaptic responses in young rats showed no such increase in old rats (**Figure 5 A - C**). Together, these observations indicate that while the direct HC drive onto the mPFC is intact, the subsequent recruitment of local mPFC circuits is diminished with age.

Reduced polysynaptic activity may arise from anatomical changes within the aged mPFC. Decreased PFC volume has been observed in aged rodents (Alexander et al., 2011), non-human primates (Alexander et al., 2008; Shamy et al., 2011), and humans (Raz et al., 2005; Salat et al., 2009; Thambisetty et al., 2010; Fjell et al., 2013) along with reductions in mPFC white-matter tracts (Makris et al., 2007; Marner et al., 2003; Pietrasik et al., 2023; Salat et al., 2005) and loss of synapses and spine density (Bloss et al., 2011, 2013). Neuron counts, however, are preserved in the mPFC between age groups across species (Haug et al., 1981; Freeman et al., 2008; Smith et al., 2004; Stranahan et al., 2012; Peters, 1998) suggesting that these changes reflect a loss of mPFC connectivity. Together these changes may impact the reliable and coordinated recruitment of polysynaptic responses to hippocampal input in the aged mPFC.

Several physiological mechanisms may also contribute to diminished neural recruitment and evoked firing rates in old rats. For example, the intrinsic excitability of aged mPFC neurons is reduced *in vitro* in rodents (Kaczorowski et al., 2012) and under anesthesia in primates (Luebke et al., 2004). In awake animals, mPFC working-memory delay activity is reduced with age (Narayanan & Laubach, 2009; M. Wang et al., 2011). Additionally, GABA_B_ receptors on parvalbumin (PV) interneurons and CB_1_ receptors on cholecystokinin (CCK) interneurons decrease with age in rodents (Albayram et al., 2012; Carpenter et al., 2016; Ginsburg & Hensler, 2022). As these receptors suppress GABA release, this loss likely increases the net inhibition onto mPFC pyramidal cells (Anastasiades et al., 2018; Carpenter et al., 2016; Liu et al., 2020), which may preferentially suppress polysynaptic excitatory responses as observed here.

### Aging has a greater impact on ventral mPFC regions

Within the mPFC, the dorsal and ventral regions are thought to play distinct functional roles. Lesion studies demonstrate that the prelimbic cortex, which corresponds to the dmPFC classification used here, supports the formation and expression of conditioned behaviors, whereas the infralimbic and dorsal peduncular cortices, which correspond to the vmPFC, mediate behavioral flexibility and suppression of previously learned responses in altered contexts (Ragozzino et al., 2002, 2003; Anderson & Floresco, 2022; Howland et al., 2022; Coutureau & Killcross, 2003; Killcross & Coutureau, 2003; Botterill et al., 2024). Recent electrophysiological studies have found that neurons in both regions are involved in decision-making, suggesting that flexible behavior likely requires coordination between them (Mukherjee & Caroni, 2018; Anderson & Floresco, 2022; Diehl & Redish, 2023). In the present study, polysynaptic fEPSP amplitude increased with depth toward the vmPFC in young rats, whereas old rats showed neither this depth gradient nor the vmPFC-dmPFC difference (**Figure 6C - D**) and the age-related reduction in polysynaptic evoked firing was most pronounced in the vmPFC (**Figure 7C - D**). Reduced vmPFC polysynaptic responses to hippocampal input may contribute to specific behavioral changes observed in aged rats, such as impaired set-shifting and reversal learning, reduced extinction learning, and impaired context-dependent memory (Poe et al., 2000; Beas et al., 2013; Caetano et al., 2012; Samson et al., 2015).

### mPFC responses evoked by intermediate and ventral hippocampal inputs are differentially affected by aging

Monosynaptic fEPSPs grew with depth along the dorsoventral mPFC axis for both stimulation sites, with iHC stimulation producing little response at dorsal depths and a sharp increase ventrally, and vHC stimulation producing a more gradual increase (**Figure 6A-B**). This pattern aligns with anatomical findings that the iHC preferentially targets the vmPFC, whereas the vHC projects to both dmPFC and vmPFC (Alemán-Andrade et al., 2025).

The age-related changes in polysynaptic responses also differed between stimulation sites. The age difference across dorsoventral gradient in polysynaptic fEPSPs reached significance only with iHC stimulation (**Figure 6C**) and polysynaptic evoked firing was reduced with age in both dmPFC and vmPFC with iHC stimulation but only in the vmPFC with vHC stimulation (**Figure 7C -D**). These patterns would be consistent with distinct populations of inhibitory and excitatory mPFC neurons engaged by the two HC inputs. The iHC input is gated by a feedforward GABAergic microcircuit in the mPFC (Gabbott et al., 2005; Takita et al., 2013), whereas vHC axons project directly onto pyramidal cells and a distinct population of inhibitory interneurons (Takita et al., 2013). The distinct short-term plasticity and neuronal targets of the two inputs (Kawashima et al., 2006; Takita et al., 2013) may further explain why iHC-evoked responses were more consistently affected by age.

### Limitations and Future Directions

All experiments were conducted under anesthesia, which can alter neural dynamics and limits direct comparisons to awake and behaving animals (Yagishita et al., 2020). Isoflurane, used in the present study, can increase the latency and reduce the amplitude of evoked LFP responses relative to other anesthetics (e.g., urethane, xylazine) and to the awake state (Sorrenti et al., 2021; Michelson & Kozai, 2018). Additionally, higher doses of isoflurane (1.4 to 2.2%) can reduce the number of recruited units and the firing rate of single units compared to awake states, and the effects on evoked-response latency vary with the type of anesthesia used. It remains to be determined whether age-related differences observed under anesthesia (Barnes, 1979) generalize to awake conditions. A second limitation is that we characterized evoked excitatory responses but could not directly assess inhibition, because baseline firing under anesthesia was too low to detect robust suppression of neural activity. Inhibitory activity is critical for understanding HC-mPFC circuit function (Isaacson & Scanziani, 2011) and could contribute to the reductions in polysynaptic responses observed here. Finally, it remains possible that a larger sample would reveal age differences in the HC-mPFC connections that appeared preserved in this study. Examining this circuit during behavior will be important for linking these physiological findings to cognitive function. Together, these findings identify a selective age-related loss of local mPFC circuit recruitment by hippocampal input, most prominent in the ventral mPFC, that may contribute to impaired cognitive flexibility in aging.

## Acknowledgements

We would like to thank Meghana Warrier and Olivia Guswiler for their assistance in collecting and processing the data. We would like to thank Peggy Nolty for administrative support. This work was supported by NIH-RF1AG081767 and the McKnight Brain Research Foundation.

## Author Contributions

SS, AV, SLC & CAB designed the experiment; SS & AV conducted the research; SS & VG analyzed the data; SS wrote the first draft of the paper; SS, CAB & SLC edited and wrote the final version of the paper. All authors reviewed the final draft of the manuscript.

